# Next-generation ultrasonic recorders facilitate effective bat activity and distribution monitoring by citizen scientists

**DOI:** 10.1101/2021.09.20.461031

**Authors:** Piia Lundberg, Melissa B. Meierhofer, Ville Vasko, Miina Suutari, Ann Ojala, Annukka Vainio, Thomas M. Lilley

## Abstract

Time and budgetary resources are often a limiting factor in the collection of large-scale ecological data. If data collected by citizen scientists were comparable to data collected by researchers, it would allow for more efficient data collection over a broad geographic area. Here, we compare the quality of data on bat activity collected by citizens (high school students and teachers) and researchers. Both researchers and citizen scientists used the same comprehensive instructions when choosing study sites. We found no differences in total bat activity minutes recorded by citizens and researchers. Instead, citizen scientists collected data from a wider variety of habitats than researchers. Involvement of citizens also increased the geographical coverage of data collection, resulting in the northernmost documentation of the Nathusius’s pipistrelle so far in Finland. Therefore, bat research can benefit from the use of citizen science when participants are given precise instructions and calibrated data collection equipment. Citizen science projects also have other far-reaching benefits, increasing, for example, the scientific literacy and interest in natural sciences of citizens. Involving citizens in science projects also has the potential to enhance their willingness to conserve nature.

**Open Research Statement:** Data are not yet provided, but will uploaded Dryad upon publication.

## INTRODUCTION

The increase in human-mediated processes such as climate change and habitat loss, have inflicted incredible pressure on the Earth’s biodiversity (Bellard et al. 2012). Worst case models predict that we are entering the sixth mass extinction through accelerated modern human-induced species losses (Ceballos et al. 2015), highlighting the need for large-scale monitoring to be initiated rapidly to gain an understanding of the impacts global change has on the biota. Despite recent advances in technology facilitating such monitoring, human resources are often a limiting factor, hindering the effective collection of large-scale spatio-temporal datasets.

Bats (order Chiroptera) are able to respond to different types of disturbance because of their unique ability amongst mammals: powered flight. Some species of bats, such as *Pipistrellus kuhlii*, have benefited from disturbances such as climate change, with an estimated range expansion of just under 400% within the last four decades (Ancillotto et al. 2016). However, populations of many species are being negatively affected by anthropogenic change in some parts or across their entire distribution range (Frick et al. 2010, Gaultier et al. 2020, Rydell et al. 2020). Consequently, growing interests lie with acquiring reliable data for the conservation and management of bats due to their importance to biodiversity, as well as their ecological and economical importance (Boyles et al. 2011, Kasso and Balakrishnan 2013).

Although monitoring methods have provided valuable information on population sizes and trends of bats over the last decades (Flaquer et al. 2007, Roche et al. 2011), these efforts have relied on manual operation of ultrasound detectors and real-time identification of bat species present. Fortunately, bat research has taken great strides forward in the last decade with the development of ultrasonic recorders and associated automated identification software (Hill et al. 2018). The use of relatively low-cost units permits the initiation of large-scale monitoring efforts producing vast amounts of data. However, researchers are still required to travel extensively for maintenance and retrieval of the units, increasing project costs.

Here, citizen science, which is a practice of engaging the public in a scientific project (McKinley et al. 2017), can assist researchers by providing spatio-temporal coverage to facilitate data collection (Devictor et al. 2010). Our project relies on the Heigl et al. (2019) description of the citizen science project, according to which the project is carried out in collaboration with citizens and researchers, it adheres to scientific standards and ethics, and relies on the flow of information between people involved in the project and transparency, including open access to data and results. Our citizen science project produces presence/absence data, which is often the case when data collection is structured and takes place under the control of researchers (Welvaert and Caley 2016). Although the quality of data collected by volunteers has come under criticism (Conrad and Hilchey 2011, Steger et al. 2017), several recent publications have shown that unpaid volunteers can produce datasets for diverse types of citizen-science projects at an accuracy that can even surpass that of professionals (Galloway et al. 2006, Kosmala et al. 2016, Brown and Williams 2019).

With the recent technological advances in automated bat recorders and species identification, the emphasis shifts towards the selection of the study site for detection of the presence or absence of given species of bats. In this study, we combined the efforts of citizen scientists and researchers to monitor the spatio-temporal distribution of two focal species: the Northern bat (*Eptesicus nilssonii*) and the Nathusius’s pipistrelle (*Pipistrellus nathusii*). We compared study site selection and total recorded bat activity between experienced bat researchers and citizens. Both species are distinguishable from all other bat species using automated identification software with manual confirmation. Whereas the Northern bat is still rather common in Finland, the Nathusius’s pipistrelle is poorly documented in Finland (Ijäs et al. 2017, Tidenberg et al. 2019, Blomberg et al. 2020). We hypothesised that given the equipment and precise instructions on how to select a study site, citizen scientists would be able to collect a comparable amount of bat activity minutes of each species as researchers.

## MATERIAL AND METHODS

### Research project participants

We reached out to the participants for our citizen science project by publishing an article about the project in Natura, a magazine published by the biology and geography teachers’ association, and by contacting the high schools directly. The biology and geography teachers enrolled their students in the project and also acted as a liaison between the school group and the researchers. The project involved students and teachers from a total of 18 schools across Finland, from Helsinki in the South to the Oulu region approximately 600km to the North. A total of 100 students (age 16-17) as well as some of the teachers participated. The number of participants from different schools varied from a single student to the entire class. From here on, we will collectively call the high school students and the participating teachers ‘citizen scientists’ for the sake of clarity. In addition to citizen scientists, three researchers collected data independently.

### Data collection

Prior to the start of the data collection, we sent the teachers the materials for two introductory lectures targeted at citizen scientists, the first on the ecology of bats and the second on how to collect data, including information on what habitat types should be selected as study sites. In addition, detailed instructions were available on our project website throughout the data collection period. We also reminded the citizen scientists about the data collection before each device deployment on the project’s Instagram account (see Online Appendix for an example). All of this was done to standardize data collection across all participants of this project. The researchers were also available throughout the data collection period for possible additional instructions and to solve problem situations.

We allowed the citizen scientists at each school to collect data alone or in small groups, due to variation in group size and preferences between schools. The most common method was to work in small groups. Once the class was divided into small groups, each small group generally had one device with which they collected data. Turns were taken within the group to deploy the device to their selected study site. Citizen scientists participating alone were allowed one or more devices as they wished. Based on the instructions given, the citizen scientists and researchers selected a study site for their device(s), which remained the same throughout the data collection period.

Researchers and citizen scientists recorded bat acoustic data using AudioMoth acoustic loggers (https://www.openacousticdevices.info/audiomoth) from 31 May–22 September 2020 (active period). We pre-programmed devices to record 10 minutes at 30-minute intervals between 21:30 and 06:00, totalling 18 recordings per night. Data were collected on two nights, from Sunday evening to Tuesday morning, every two weeks. All devices initiated recording automatically at 21:30 and recorded according to the same schedule for two nights. Citizen scientists and researchers deployed the devices at their study sites prior to each data collection period and retrieved the devices from their study sites after the two-day data collection.

A total of 324 10-minute recordings could be collected by one device during the data collection period. Sometimes the recording failed (e.g., due to the device getting wet), resulting in missing data. Altogether, we used 121 AudioMoths in the data collection, of which 52 were used by the citizen scientists and 69 by three researchers. We did not receive location data for two devices, reducing the total number of devices used in this study to 119 (Figure 1).

**Figure 1.**
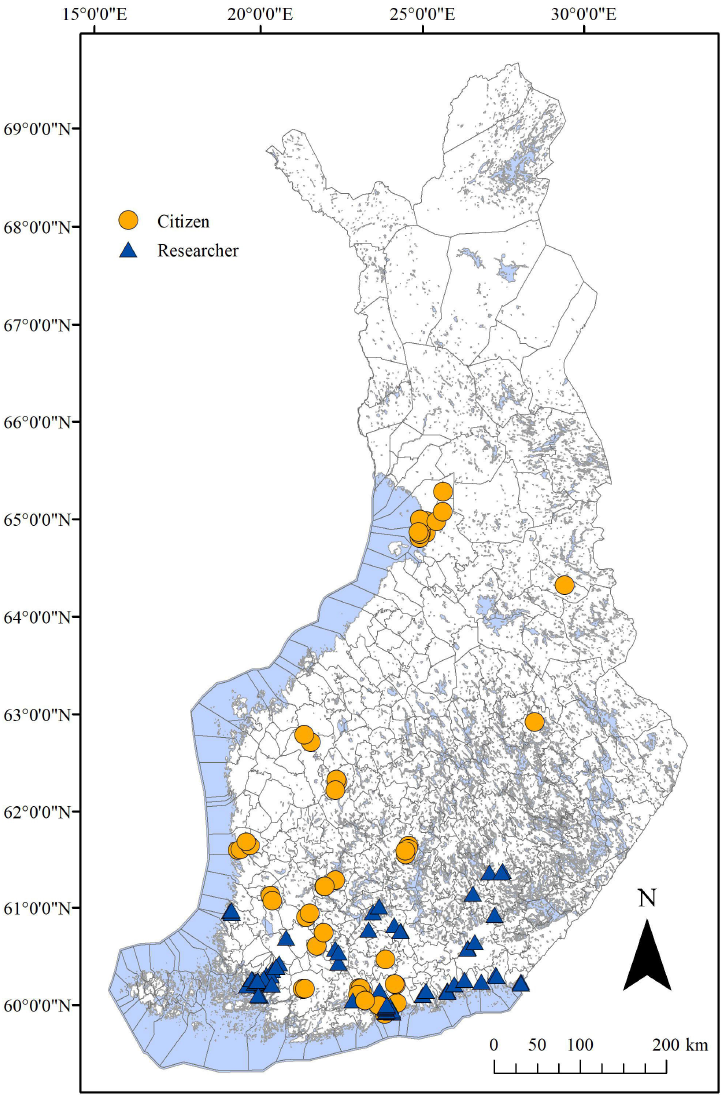
Map of the distribution of deployed AudioMoths (n = 119) across Finland by citizens (circle, n = 50) and researchers (triangle; n = 69).

In addition, citizen scientists and researchers added the location of their study site to the Finnish Biodiversity Information Facility (FinBIF), an online data depository maintained by the Finnish Museum of Natural History, https://laji.fi/en, on which a separate form (Lepakkolomake,“a bat form”) had been created for this study. Citizen scientists and researchers added the following information to the bat form: a description of the habitat of the study site, environmental data (including rainfall, temperature and relative humidity) related to their study sites. We had a list of 16 habitat variables (e.g., sparse forest, lake shore, edge of shore coppice or yard), and the participants selected one or more variables that described their study site. In addition, they were asked to upload photos of the surroundings of their study site to the online form. However, because our study focuses on comparing the total activity minutes of bats recorded by citizens and researchers, we only included habitat type from the environmental data gathered in our statistical analyses.

### Data analysis methods

#### Identification of bat species using Kaleidoscope Pro

Identification of bat species was conducted with Kaleidoscope Pro (Wildlife Acoustics, Inc., v. 5.1.9). We used AutoID for bats, split the data into 15-second WAV-files and deleted noise files. The following settings were used in the analysis: signal detection parameters were set with frequency between 8 and 120 (kHz), minimum and maximum length of detected pulses 1–200 (ms), minimum number of pulses 1 and the maximum inter-syllable gap of 500 (ms). To effectively locate all incidences of *E. nilssonii* and *P. nathusii* in the data set, we also searched for additional species, which are sometimes calls of our focal species incorrectly sorted by the algorithm. These additional species were *E. serotinus*, *Myotis dasycneme*, *Nyctalus leisleri*, *P. pipistrellus*, *P. kuhlii*, and *Vespertilio murinus* (see Online Appendix for more detailed description). The software saves the post-processed outputs as csv-files with their associated WAV-files, which we then checked manually as misidentification by software can occur (Rydell et al. 2017).

We then combined the separate csv-files into one dataset and added the following information: location of the device (i.e., latitude), individual ID (i.e., who collected the data), and observer type (i.e., citizen or researcher). In addition, we classified the habitats of the study sites into three categories: 1) coastal areas and wetlands, 2) sparsely wooded areas and forests and 3) open landscape (courtyards, parks, agricultural areas). When classifying the habitats, we used descriptions and any photos of the study site provided by the participants, topographic maps, freely available aerial photographs from Finland. Finally, we calculated bat activity minutes: If any of the four 15-second files within a minute contained a focal bat species, the minute was tagged as an active minute. The sum of active minutes per site was used as the response variable.

#### Statistical analyses

We used a linear mixed effects model (function glmer in package lme4; Bates et al. 2015) with activity minutes (positive outcomes) of the total recording minutes (total trials) as the response variable, latitude and observer as fixed effects, and individual ID as a random effect, to test whether citizens or researchers were better at collecting acoustic data for the Northern bat. We fitted the model with a binomial distribution. Unfortunately, we had to omit enough Nathusius’s pipistrelle from our analyses because we did not have enough activity data to answer our question. To determine whether environments of sites chosen by researchers and citizens differed, we conducted a Fisher’s exact test. We considered a *p*-value ≤ 0.05 significant for all tests. We used R v. 3.5.0 to conduct all analyses.

## RESULTS

Citizens and researchers recorded for a total of 148,530 and 197,810 minutes, respectively. We had only one site without bat activity. Nathusius’s pipistrelle were present at 31 out of 119 sites and Northern bats at 118 out of 119 sites. Of the total recorded minutes, citizens and researchers recorded a total 4853 and 14276 minutes of Northern bat activity, respectively, and 40 and 603 minutes of the Nathusius’s pipistrelle, respectively (Table 1). Of these 603 minutes, 89.9% were from recordings of three devices (of which the best device accounted for 69.3% of all minutes) (Table 1). Citizens deployed devices across all latitudes from 60 to 65 degrees N, whereas researchers deployed devices between 60 and 61degrees N (Figure 1). Northern bat activity increased with decreasing latitude (*β* = −0.53, z = −20.15, *p* < 0.05; Figure 2A). Although there was no significant difference between researchers and citizen scientists in detecting Northern bats (*β*= 0.45, z = 1.46, *p* = 0.14), researchers tended to place devices in locations where more Northern bat activity was recorded (Figure 2B). There was, however, a significant difference between researchers and citizens on selection of habitat type for acoustic logger deployment (*p* < 0.05).

**Table 1.** Total recording time and total bat activity for the northern bat (*E. nilssonii*) and Nathusius’s pipistrelle (*P. nathusii*) across latitudes monitored by observer type (citizen or researcher).

|  | Total Recording<br>Minutes | Total northern bat<br>Activity Minutes | Total Nathusius'<br>pipistrelle Activity<br>Minutes | Total<br>devices |
| --- | --- | --- | --- | --- |
| Citizen | 4893 | 4853 | 40 | 50 |
| 60° | 3057 | 3029 | 28 | 18 |
| 61° | 1223 | 1212 | 11 | 15 |
| 62° | 411 | 411 | 0 | 4 |
| 63° | 109 | 109 | 0 | 2 |
| 64° | 4 | 4 | 0 | 1 |
| 65° | 89 | 88 | 1 | 10 |
| Researcher | 14879 | 14276 | 603 | 69 |
| 60° | 12473 | 11874 | 599 | 55 |
| 61° | 2406 | 2402 | 4 | 14 |

**Figure 2.**
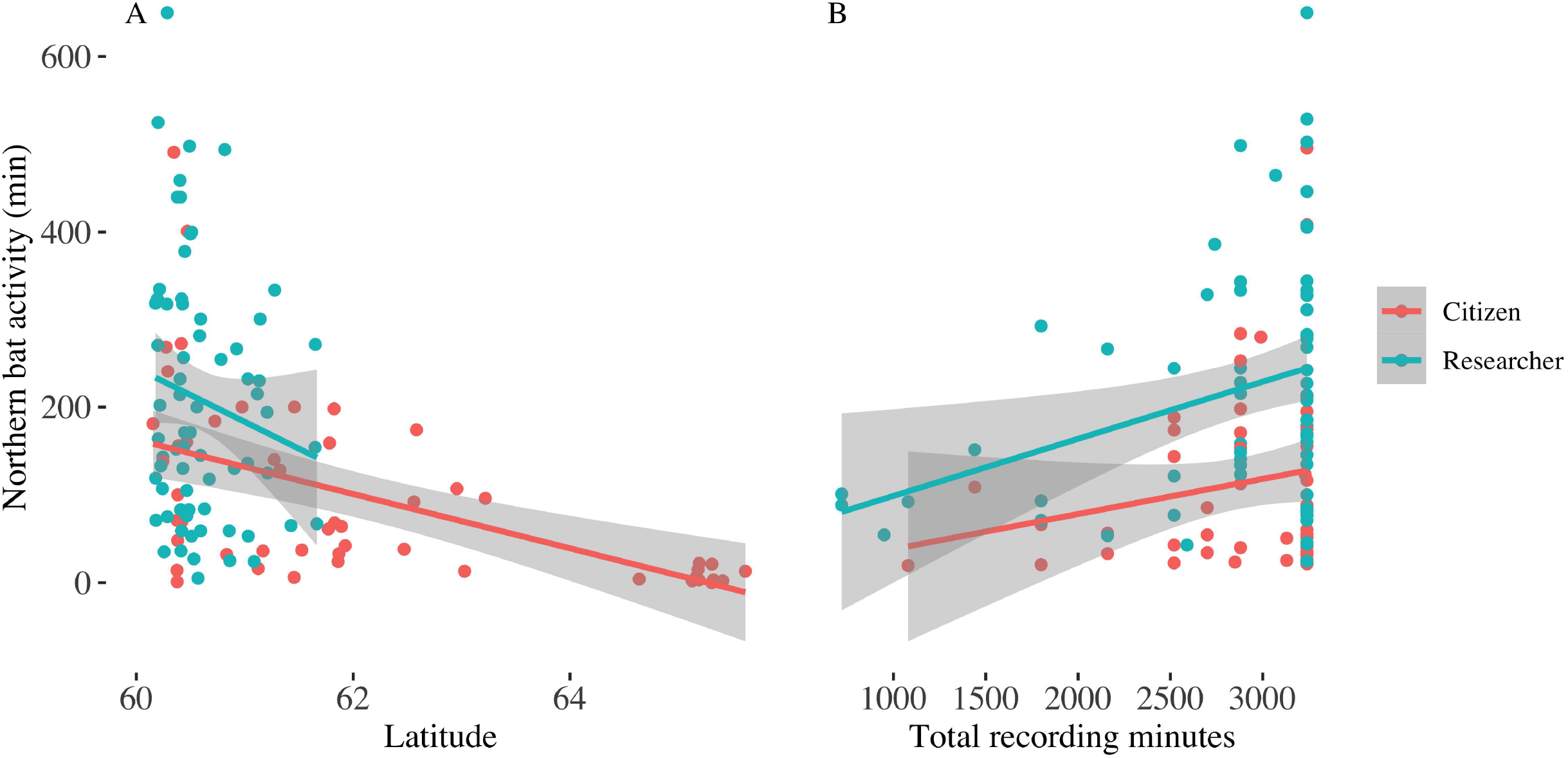
Northern bat (*E. nilssonii*) activity recorded across latitudes (A) and based on total recording minutes (B). Gray region represents the 95% confidence level interval for predictions from a linear model.

## DISCUSSION

Citizen volunteers can help researchers by collecting large amounts of data, but the quality of the data gathered by citizen scientists remains to be determined (Wiggins et al. 2011, Kosmala et al. 2016). Our study addressed this issue by investigating the use of citizen science in studying bat activity in Finland with next-generation ultrasonic recorders. Although researchers recorded more data, we found no significant difference between citizen scientists and researchers in recorded bat activity minutes. However, we acknowledge that clustering of sampling locations at some latitudes may have affected the results in this comparison. In our case, citizen science proved an effective method: it provided data that was as good as the data produced by researchers, but from a much larger geographical area. In a similar way, citizen science has produced reliable information on the distribution of birds (Fournier et al. 2017, Biddle et al. 2021). Structured citizen science projects often bring together large numbers of people at the same time in the same place, which reduces their spatio-temporal coverage, compared to, for example, crowdsourcing-type research (Welvaert and Caley 2016). Our project, however, differs from this in that we collaborated with high schools remotely thereby increasing our regional coverage.

Our study focused on two species of bats with differing population trajectories: the Nathusius’s pipistrelle and the Northern bat. The former is a species that is rapidly expanding its range to the north in Europe (Lundy et al. 2010, Blomberg et al. 2020), whereas the latter, although still abundant, has shown a sharp decline in population size (Rydell et al. 2020). However, both of the species have rather broad habitat preferences (Tidenberg et al. 2019), and had we chosen to focus on species with more specific habitat requirements, we may have seen differences between the two groups of observers in collected bat activity due to placement of the recorders. Our study suggests that citizen science also has potential in studying the occurrence of a species with insufficient data especially by covering a broader geographical area than researchers alone. With the help of citizen scientists, we produced valuable occurrence information for the Nathusius’s pipistrelle, with the current northernmost observation in Finland in the Oulu region (c. 65° north) recorded by participating students. Other citizen science projects have produced new information in a similar way, for instance, on the distribution of rare fish species (Naasan Aga Spyridopoulou et al. 2020, Tiralongo et al. 2020), insects (Zapponi et al. 2017, Soroye et al. 2018, Wilson et al. 2020) and large carnivores in remote areas (Farhadinia et al. 2018).

There are still unanswered questions on the reliability of data collected by non-professionals that need to be accounted for when implementing citizen science projects. Citizen science may be subject to biases depending on the sampling design (Geldmann et al. 2016, Brown and Williams 2019) or through interpretations made by citizens themselves (Galloway et al. 2006), which has been noted, for instance, in the over-reporting of rare species (Galloway et al. 2006, Gardiner et al. 2012). Acoustic monitoring does not require interpretation (or identification of the species) from participants, making it less susceptible to such bias, which also supports its use in bat-orientated citizen science projects.

Another potential source of bias is the choice of study sites (Dambly et al. 2021), which necessitates the need for standardization of data collection. We provided detailed instructions for site selection and data collection to maintain consistency across all participants. We also pre-programmed the entire recording schedule on the devices to ensure ease of data collection for the participants as well as simultaneous data collection at all study sites. Training of the participants, which is common in structured citizen science projects, was carried out in our project with the help of biology and geography teachers both having a core understanding of ecological principles. This, together with the commitment and active involvement of the participants, enables the production of high-quality data for the use of researchers (Welvaert and Caley 2016). Consequently, potential uncertainties associated with the use of citizen science (e.g. Kosmala et al. 2016, Steger et al. 2017, Brown and Williams 2019), can be mitigated through means of careful planning of the study (Kosmala et al. 2016) and designing the research protocol from the perspective of citizen scientist involvement (Cohn 2008). Nevertheless, we recommend that projects using citizen science carefully consider appropriate study design and methods of data collection and invest in instructing the participants.

In addition to enabling the collection of data that would not be otherwise possible (Chandler et al. 2017), the benefits of citizen science extend beyond science. For example, citizen science enables a bidirectional information flow between researchers and the public, and increases scientific and environmental literacy among the participants (Trumbull et al. 2000, McKinley et al. 2017). Furthermore, collecting environmental data has also been found to foster engagement in environmental conservation actions among volunteers, increasing community interaction and interest toward conservation (Ballard et al. 2017). The possibility of being actively in contact with nature cannot be underestimated in modern urbanised societies. Engagement in nature activities and positive nature experiences are associated with higher felt connection with nature, willingness to take care of nature, and even with higher personal subjective well-being and happiness (Zelenski and Nisbet 2014, Martin et al. 2020, Cleary et al. 2020).

For schools citizen science provides an opportunity for students to learn more about science (Trumbull et al. 2000) and learning outside the classroom (Hulbert 2016). As a part of this project, we designed a customized science course on bats for the high schools that participated in our project including all stages of scientific research. The participants of the course had the opportunity to analyse the data they collected using a free version of Kaleidoscope or Audacity software to manually identify the bats according to the tutorial we had prepared for them. These data were not used in our analyses. The purpose of this course was to increase the understanding of the scientific process through practical assignments. At the end of the course, we held an online lecture that summarized the results of the project. We also shared information about bats through the project’s Instagram account, along with additional instructions.

Currently, citizen science is included in only a small proportion of academic research publications (Callaghan et al. 2021). One of the main barriers for use of citizen science has been the question on data quality. The findings of this study indicate that volunteers can collect high quality data using novel digital innovations, when given good instructions. Another barrier is the lack of legitimacy of citizen science in scientific communities (Burgess et al. 2017, Golumbic et al. 2017). Therefore, more research focusing on the quality of data gathered by citizen scientists is needed to make greater use of the potential of citizen science in scientific research.

## CONCLUSIONS

Citizen science provides numerous invaluable opportunities in understanding the effects of e.g., biodiversity loss and climate change through environmental monitoring, thus stressing the need to enhance the use of citizen science in research. However, careful consideration should be given to study design and to the instructions provided to citizen scientists to ensure the quality of the data. The main advantage of citizen science to the field of research is the broad geographical coverage, which could not be achieved by researchers alone due to schedule and budgetary reasons. Another advantage is the wide range of habitats covered by citizen scientists. Citizen science can also be useful in studying the occurrence of rare species. Furthermore, citizen science projects have additional benefits, such as increasing the knowledge of the participants, and interest in nature and natural sciences.

## ACKNOWLEDGEMENTS

We thank our citizen science project participants for their help in data collection, without whom this study would not have been possible, as well as the Kone Foundation for funding this project (Funding ID: 201802425, Natural Resources Institute Finland (Luke) Project nr: 41007-00165400). We thank Lasse Ruokolainen for his advice in statistical analysis and Janne Lundberg for his help with pre-programming the AudioMoth devices for the data collection. We also thank Finnish Museum of Natural History ICT-team and CSC – IT Centre for Science for assistance in data handling.

