## Appendix file for "Next-generation ultrasonic recorders facilitate effective bat activity and distribution monitoring by citizen scientists"

Figure 1. An example of our Instagram post where we remind (and instruct) our citizen scientists on how to deploy the AudioMoth devices

Detailed description of the methods

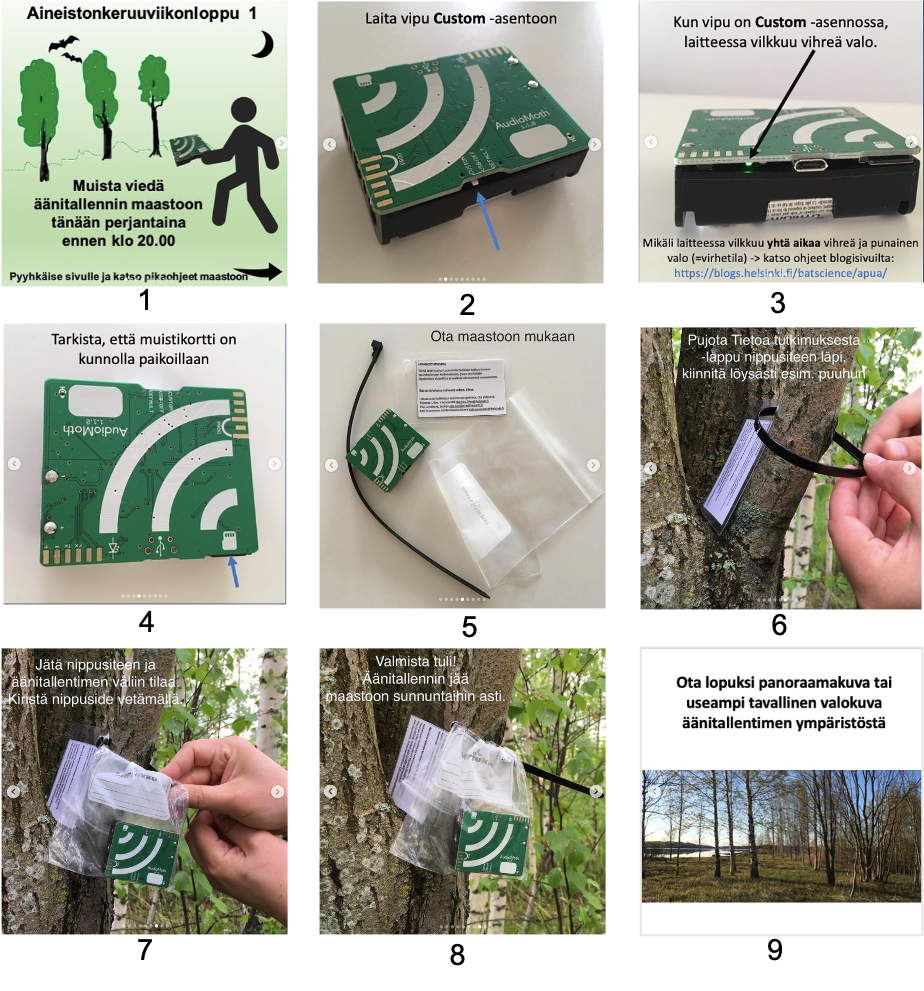

**Figure 1.** Instructions to students for collecting audio data. The instructions were distributed through the project Instagram-account over the study weekends. 1) Remember to take the audio recorder to your study location on Friday before 10PM. 2) Set the switch to the custom-position. 3) A green light will flash when the switch is in the custom-position. In case both the red and green light flash simultaneously (error mode) -> see instruction on the blog-page. 4) Ensure that the memory card is inserted correctly. 5) Take these with you to the study location. 6) Thread the cable tie through the hole on the information card and secure cable tie loosely around a tree branch or similar. 7) Leave some room between the cable tie and the audio recorder. Tighten the cable tie. 8) You’re set! The audio recorded will remain at the location until Sunday. 9) Finally, take a panorama photo of the recorder and the surroundings.

**DETAILED DESCRIPTION OF THE METHODS**

For calls marked as Enil, we checked only the weakest pulses with a matching value of 2 or less. As Enil can be misidentified as Eser, Mdas, Mbec, Nlei and Vmur, we checked all calls marked as the aforementioned species manually. Similarly, all Pnat, Pkuh, Ppip and Ppyg files were checked to determine whether or not they were Pnat as the software can misidentify these species. We checked all Paur and Paus files for Enil and Paur. In addition to the AutoID, the software provides one or two alternate species in cases when the call is not clear or multiple species are present in the same file. We checked all files where Enil or Pnat were suggested as alternate species. Finally, we went through all files specified as NoID and manually identified the species if present.
